# Genus-specific remodeling of carbon and energy metabolism facilitates acetoclastic methanogenesis in *Methanosarcina* spp. and *Methanothrix* spp.

**DOI:** 10.1101/2025.10.01.679765

**Authors:** Blake E. Downing, Dinesh Gupta, Katie E. Shalvarjian, Dipti D. Nayak

## Abstract

Methanogenic archaea (methanogens) are microorganisms that obligately produce methane as a byproduct of their energy metabolism. While most methanogens grow on CO_2_+H_2_, isolates of the Genus *Methanosarcina* and *Methanothrix* can use acetate as the sole substrate for methanogenesis. Methanogenic growth on acetate, i.e., acetoclastic methanogenesis, is hypothesized to require two distinct genetic modules: one for the activation of acetate to acetyl-CoA and the other for producing a chemiosmotic gradient using electrons derived from ferredoxin. In *Methanosarcina* spp., the activation of acetate to acetyl-CoA is mediated by acetate kinase (Ack) and phosphotransacetylase (Pta) whereas *Methanothrix* spp. encode AMP-forming acetyl-CoA synthetases (Acs). The *Rhodobacter* nitrogen fixation complex (Rnf) or Energy converting hydrogenase (Ech) are critical for energy conservation in *Methanosarcina* spp. during growth on acetate, and a F_420_:phenazine oxidoreductase-like complex (Fpo’) likely plays an analogous role in *Methanothrix* spp. Here, we tested the proposed modularity of these pathways to facilitate acetoclastic methanogenesis. First, we surveyed over a hundred genomes within the Class *Methanosarcinia* to show that the genomic potential for acetoclastic methanogenesis using distinct combinations of modules is widespread. We then used the genetically tractable strain, *Methanosarcina acetivorans,* to build all modular combinations for acetoclastic methanogenesis. Our results indicate that Acs, while functional, cannot replace Ack+Pta to rescue acetate growth in *M. acetivorans*. Similarly, the Fpo’ bioenergetic complex cannot replace Rnf. As such, our work suggests that, in addition to horizontal gene transfer of core catabolic modules, acetoclastic metabolism in methanogens requires changes in energetic modules too.

**IMPORTANCE:** A large fraction of biogenic methane is derived from acetate, yet acetoclastic methanogens, i.e., methanogens that grow on acetate, remain poorly characterized due to their slow growth. Two groups of methanogens, *Methanosarcina* spp. and *Methanothrix* spp., perform acetoclastic methanogenesis using distinct sets of genes for acetate activation and energy conservation. It is widely hypothesized that these genetic modules from *Methanosarcina* spp. and *Methanothrix* spp. are functionally analogous and would thus be interchangeable. To test this hypothesis, we engineered different combinations of modules for acetoclastic growth in *Methanosarcina acetivorans*. Our results challenge this hypothesized paradigm of modularity, and we posit that other changes to the carbon and electron transfer pathways were crucial for the emergence of acetoclastic methanogenesis.

## INTRODUCTION

Methane (CH_4_) is a potent greenhouse gas that traps heat in Earth’s atmosphere up to one hundred times more effectively than an equivalent amount of carbon dioxide (CO_2_) over a period of twenty years (1, 2). An accurate accounting of the sources and sinks of methane is important for modeling Earth’s climate in the past, present, and future. A large fraction of biogenic methane released into the atmosphere is produced by methanogens, microorganisms that generate methane as a by-product of their energy metabolism (1, 3). Methanogens are ubiquitous in anoxic environments ranging from sediments to the human gastrointestinal tract (3), and their growth substrates range from inorganic (H_2_ + CO_2_ and formate) to organic (C_1_ compounds or acetate) depending on the environment (3–5). In contemporary times, acetate often fuels methanogenesis in human-built environments like waste-water treatment facilities and landfills, but this process evolved at least 250-million years ago when methanogens belonging to the Genus *Methanosarcina* acquired genes for acetate catabolism by horizontal gene transfer from bacteria (5, 6).

Acetoclastic methanogenesis has been demonstrated in two distinct Genera within the Class *Methanosarcinia*: *Methanosarcina* and *Methanothrix* (5, 7). *Methanosarcina* spp. are metabolic generalists with a broad substrate range that includes H_2_+CO_2_, C_1_ compounds like methanol or methylamines, and acetate (5, 7). In contrast, *Methanothrix* spp. are metabolic specialists that can only grow on acetate (5, 7). A growth rate versus yield tradeoff likely allows these two groups of methanogens to occupy distinct ecological niches (5). *Methanosarcina* spp. have faster growth rates and are typically found in acetate-rich environments, whereas *Methanothrix* spp. have a higher substrate affinity for acetate and thrive in acetate-limited regimes (5). These ecological distinctions are thought to stem from different pathways for activating acetate to acetyl-coA and energy conservation modules in the two groups of methanogens (**Fig. 1A**) (5). In the Genus *Methanosarcina*, acetate activation proceeds through the combined activities of two enzymes, acetate kinase (Ack) and phosphotransacetylase (Pta), where Ack hydrolyzes an ATP to activate acetate to acetyl-phosphate and Pta catalyzes the transfer of the acetyl-group to coenzyme A (CoA) to produce acetyl-CoA (**Fig. 1B**) (5). In the Genus *Methanothrix*, acetate activation to acetyl-CoA occurs in one step via the AMP-forming acetyl-CoA synthetase (Acs) (5). The regeneration of ATP from AMP requires the hydrolysis of a second molecule of ATP as shown in (**Fig. 1B**) (5). Hence, the Acs-dependent acetate activation pathway in *Methanothrix* spp. requires twice as much energy investment than the Ack + Pta activation pathway in *Methanosarcina* spp. The enzymes involved in the dismutation of acetyl-CoA to produce CO_2_ and methane are largely the same and have been reviewed elsewhere (**Fig. 1A**) (4, 5, 7). During acetoclastic methanogenesis, electrons are generated in the form of reduced Ferredoxin (Fd_red_), which are used to generate a chemiosmotic gradient for ATP synthesis via an electron transport chain (ETC) (3, 8). *Methanosarcina* spp. use either the *<u>R</u>hodobacter* <u>n</u>itrogen fixation (Rnf) complex or the membrane-bound energy-<u>c</u>onverting <u>h</u>ydrogenase (Ech) to generate a Na^+^ or a H^+^ gradient with electrons derived from Fd_red_, respectively (**Fig. 1C**) (8–13). Owing to their slow growth and genetic intractability, the ETC of *Methanothrix* spp. is not well-resolved (14). However, biochemical assays with crude membrane preparations and genomic sequencing suggest that a modified form of the <u>F</u>_420_:<u>p</u>henazine <u>o</u>xidoreductase (Fpo) lacking a F_420_-interacting “head” subunit, FpoF, (Fpo’) uses the electrons derived from Fd_red_ to generate a proton gradient (**Fig. 1C**) (15).

**Figure 1.**
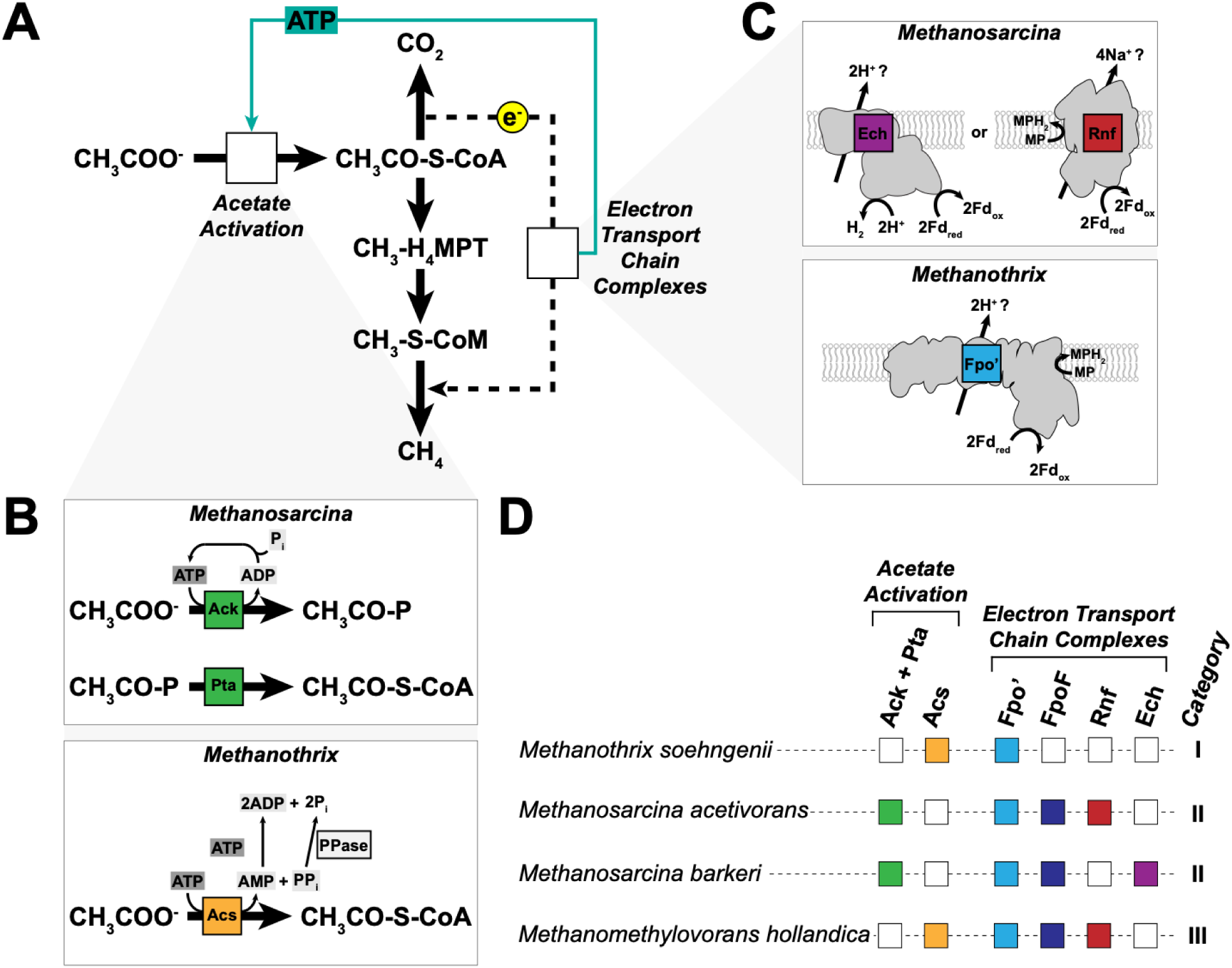
Pathways for acetate consumption and energy conservation across the Class *Methanosarcinia*. (**A**) Pathways of carbon (solid arrows) and electron (dashed arrow) flow during acetoclastic methanogenesis. Acetate is first activated in an ATP-dependent manner to acetyl-CoA, which is dismutated into its methyl and carbonyl groups. The methyl group enters the methanogenesis pathway via the C1-carrier tetrahydromethanopterin (H_4_MPT) [or a derivative in select methanogens, tetrahydrosarcinopterin (H_4_SPT)] and is subsequently transferred to another C1 carrier, coenzyme M (CoM). The carbonyl group is oxidized to carbon dioxide and reducing equivalents from the oxidation (e^-^) pass through the electron transport chain. The electron transport chain provides the electrons necessary for the reduction of the methyl group to methane, while simultaneously generating the ATP necessary for substrate activation (solid teal arrow). For simplicity, the carbon and electron flow steps do not show cofactors and/or electron carriers as substrates/products. Empty boxes labeled “Acetate Activation” and ‘Electron Transport Chain Complexes” note where proteins catalyzing these functions differ between methanogens. (**B**) Schematics of acetate activation in different acetoclastic methanogens. In *Methanosarcina* spp., acetate is activated in an ATP-dependent manner to acetyl-phosphate (acetyl-P) by acetate kinase (Ack), which is then converted to acetyl-CoA through the activity of a second enzyme, phosphotransacetylase (Pta). The single ATP spent during activation is regenerated to ATP likely through the activity of ATP synthase or other ADP-dependent enzymes. In *Methnothrix* spp., acetate is activated directly to acetyl-CoA in a single, ATP-dependent step catalyzed by acetyl-CoA synthetase (Acs) coupled to the production of AMP and pyrophosphate. AMP is then regenerated to ADP through the hydrolysis of a second equivalent of ATP. The pyrophosphate is hydrolyzed by a soluble pyrophosphatase (PPase) and can be used to regenerate the two equivalents of ADP to two ATP. (**C**) Schematics of the electron transport chain complexes involved in energy conservation during acetoclastic methanogens. In *Methanosarcina* spp., reduced ferredoxin generated by acetoclastic methanogenesis (Fd_red_) is re-oxidized via either the proton-translocating Ech (energy converting hydrogenase) complex coupled to hydrogen gas (H_2_) production, or the sodium-translocating Rnf (*Rhodobacter* nitrogen fixation) complex coupled to methanophenazine (MP) reduction (MPH_2_). In *Methanothrix*, re-oxidation of Fd_red_ coupled to MP reduction is thought to be catalyzed by a proton-translocating F_420_:phenazine oxidoreductase homolog that lacks the coenzyme-F420 active site-containing subunit (Fpo’). (**D**) Presence/absence of acetate activation and electron transport modules in four representative methanogen species across the *Methanosarcinia*. Categories “Acetate Activation” and “Electron Transport Complexes” refer to the genes shown in parts A and B, respectively. For each species, the presence or absence of the protein(s) is indicated by a color-filled or empty box, respectively: Ack + Pta (acetate kinase + phosphotransacetylase), green; Acs (acetyl-CoA synthetase [AMP-forming]), yellow; Fpo’ (F_420_:phenazine oxidoreductase lacking FpoF subunit), light blue; FpoF (coenzyme F_420_ active site-containing subunit of F_420_:phenazine oxidoreductase), dark blue; Rnf (*Rhodobacter* nitrogen fixation complex), red; Ech (energy converting hydrogenase), purple.

Despite their overwhelming contribution to the global methane budget, physiological studies with acetoclastic methanogens, especially *Methanothrix* spp., have been sparse, especially in the last decade (16). Here, we test if the distinct pathways for acetoclastic methanogenesis from *Methanosarcina* spp. and *Methanothrix* spp. are modular and cross-compatible. First, we surveyed sequenced methanogen genomes to identify strains that might have the potential to perform acetoclastic methanogenesis using a profile hidden Markov model (HMM)-based screen. While the specific combination of acetate activation genes and energy conservation modules found in *Methanothrix* spp. or *Methanosarcina* spp. are absent in other genomes, alternate combinations, especially Acs and Rnf, are more broadly distributed. We then engineered strains of *Methanosarcina acetivorans* with different combinations of activation pathways (Ack+Pta versus Acs) and bioenergetic modules (Rnf/Ech versus Fpo’) that might fuel acetoclastic methanogenesis. Our findings suggest that existing combinations of acetate activation and energy conservation are intricately linked, such that these two modules must co-evolve to facilitate methanogenic growth on acetate.

## RESULTS

### Genomic analysis highlights the potential for novel acetoclastic methanogenesis pathways

We used a custom, profile HMM-based search tool to survey extant methanogens for the presence of proteins that play a role in either acetate activation or energy conservation during acetoclastic methanogenesis. We restricted our search to sequenced genomes available through the Genome Taxonomy Database (GTDB r214.0) within the Class *Methanosarcinia* (n=133 genomes) as it comprises most known methanogens with an ETC, which is essential for energy conservation during acetoclastic growth.

First, we surveyed proteins involved in acetate activation, either Ack + Pta or Acs. We found that Ack + Pta is only present in *Methanosarcina* spp. (94%, n=29/31, **Fig. S1, Table S1**) and in some members of the Genus *Methanimicrococcus* (50%, n=3/6), including the type-strain *Methanimicrococcus blatticola* (17). In contrast, Acs is more broadly distributed as we found hits within *Methanothrix* (89%, n=24/27), *Methermicoccus* (100%, n=1), *Methanomethylovorans* (86%, n=6/7) and multiple other Genera (**Fig. S1, Table S1**). We did not detect the co-occurrence of Ack + Pta and Acs in any of the sequenced genomes.

Next, we searched for energy conservation modules that can use Fd_red_ to generate an chemiosmotic gradient, i.e., Rnf, Ech, or Fpo’ (8, 12). Rnf is broadly distributed in several different Genera including *Methanosarcina* (26%, n=8/31), *Methanolobus* (100%, n=19/19), *Methanomethylovorans* (86%, n=6/7) (**Fig. S1, Table S1**). However, we were only able to detect Ech in members of *Methanosarcina* (71%, n=22/31, **Fig. S1, Table S1**). Since the canonical Fpo complex interacts with F_420_ via FpoF (18), we surveyed genomes for all subunits of the Fpo complex excluding FpoF (Fpo’) and separately searched for FpoF to distinguish between these potential variants. While Fpo (Fpo’ + FpoF) is broadly distributed within the Class *Methanosarcinia* (56%, n=75/133, **Fig. S1, Table S1**), genomes that solely encode Fpo’ are more limited (26%, n=34/133) and are typically restricted members of the Genus *Methanothrix* (81%, n=22/27) and *Methermicoccus* (100%, n=1, **Fig. S1, Table S1**).

Based on our survey, we developed a classification scheme to describe patterns of co-occurrence between acetate activation and energy conservation modules (**Fig. 1D**). We define the Category I genomes as those containing Acs for acetate activation and Fpo’ for energy conservation (23% of *Methanosarcinia*, n=30/133), as exemplified by *Methanothrix soehngenii*. Category II genomes include those that use Ack + Pta to activate acetate, and either Rnf or Ech to conserve energy (25% of *Methanosarcinia*, n=33/134). Most Category II genomes also encode a complete Fpo complex (82% of Category II, n=27/33). Category II is represented by genomes such as *Methanosarcina acetivorans* and *Methanosarcina barkeri* (**Fig. 1D**) We define Category III genomes as those containing the Category I acetate activation module (Acs) alongside the Category II energy conservation modules (i.e., Rnf), thus representing a hybrid between the first two categories (32% of *Methanosarcinia*, n=42/133) (**Fig. 1D**). *Methanomethylovorans hollandica* represents a Category III genome. Notably, acetoclastic methanogenesis has not yet been demonstrated in a Category III strain. Intriguingly, we did not find a single genome that encodes Ack + Pta and just Fpo’, another potential combination of modules that might facilitate acetoclastic methanogenesis.

Our analysis raises questions about the cross-compatibility between substrate activation and energy conservation modules in acetoclastic methanogens. In other words, even though the acetate activation and energy conservation modules perform the same biochemistry, are they functionally analogous and interchangeable? Further, how has the interaction between activation enzymes and respiratory complexes shaped the evolutionary history of acetoclastic methanogens? To address these questions, we harnessed *M. acetivorans* as an experimental chassis to explore if: (i) the naturally occurring combination of Acs and Rnf (as in Category III genomes) or (ii) a synthetic combination of Ack + Pta and Fpo’ can support acetoclastic methanogenesis *in vivo*.

### Acetyl-CoA synthetases are functional but do not support acetoclastic growth of *M. acetivorans*

First, we explored the possibility of acetoclastic methanogenesis in Category III strains (**Fig. 1D**). Theoretically, these methanogens could grow on acetate using Acs for acetate activation and conserve energy using the Rnf complex. To test this possibility, we chose to engineer *M. acetivorans,* a category II strain, into a Category III strain rather than test for acetate growth in a naturally occurring Category III methanogen. We opted for the former because acetate growth under laboratory conditions is well established in *M. acetivorans* whereas the latter may need unknown regulatory cues to stimulate acetate growth. As the first step in the engineering process, we deleted the native acetate activation module (Δ*ack-pta*, MA3606-3607, MA_RS18805-18810), in the parent strain (WWM60 or wildtype, WT). We verified the absence of any off-target mutations due to CRISPR-editing and the presence of an in-frame markerless chromosomal deletion of the *ack-pta* genes by whole genome sequencing (**Table S2**). To validate the absence of Ack and Pta, we also assayed for acetyl-CoA production in cell lysates using the acetylhydroxamate assay as previously described (19, 20). The mutant had *ca.* 7% of the activity observed in the WT strain, which did not seem to vary in a substrate-dependent manner (**Fig. 2A**). This mutant was also incapable of growth on acetate as the sole substrate for methanogenesis, even after prolonged incubation for 6+ months (**Fig. 2B**, **Table 1**). Curiously, despite the ability to generate acetyl-CoA on TMA, WT does not have a detectable growth advantage in minimal medium containing both TMA and acetate compared to the Δ*ack-pta* mutant (**Fig. S2, Table S3**). To further corroborate that the production of acetyl-CoA in WT is due to the activity of Ack+Pta, we complemented the Δ*ack-pta* mutant *in trans* with a plasmid expressing the *ack* and *pta* locus from a tetracycline-inducible promoter (Δ*ack-pta/PmcrB(tetO4)-ack+pta*). Acetyl-CoA production and acetate growth were restored in the complementation mutant (**Fig. 2**, **Table 1**).

**Figure 2.**
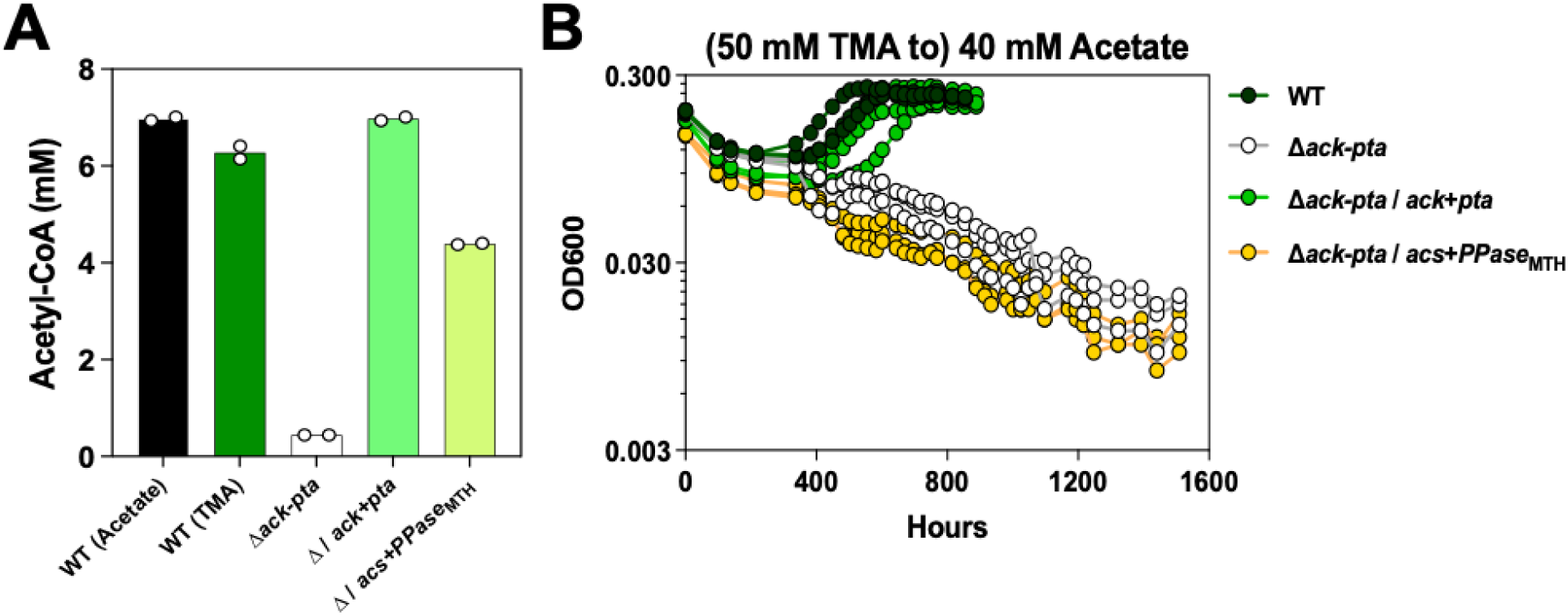
Complementation of the *M. acetivorans ack-pta* deletion mutant with *acs* does not restore growth on acetate. (**A**) Quantification of acetyl-CoA production from cell lysates using the acetylhydroxamate assay. Assays were conducted with WWM60 (wildtype or WT) grown in high-salt (HS) minimal medium with 40 mM acetate (black bar) or 50 mM TMA (trimethylamine) (dark green bar), and the Δ*ack-pta* mutant (white bar), Δ*ack-pta* expressing *ack+pta in trans* (Δ*ack-pta/PmcrB(tetO4)-ack-pta*) (light green bar), and the Δ*ack-pta* mutant expressing *acs*+*PPase*_MTH_ *in trans* (Δ*ack-pta/PmcrB(tetO4)-acs*-*PPase*_MTH_) (yellow-green bar) all grown in HS medium with 50 mM TMA and 2 µg/mL Puromycin. All reactions were performed with two biological replicates. Each replicate comprised of 1 mg total protein that was incubated at 37°C for 30 mins under the assay conditions (see Materials and Methods). (**B**) Growth curves of WT (dark green circles), the *Δack-pta* mutant (white circles), and complemented strains Δ*ack-pta*/*ack-pta* (light green circles) and *Δack-pta/acs+Ppase*_MTH_ (yellow circles). Cells were pre-grown in high-salt (HS) with 50 mM TMA and inoculated into HS medium with 40 mM acetate with or without 2 µg/mL Puromycin for maintenance of the complementation plasmid and 100 µg/mL tetracycline for full induction of the complemented gene. Three replicate growth curves were conducted for each strain. Growth parameters for each strain are shown in Table 1.

**Table 1.** Growth characteristics of the WWM60 (wildtype or WT), the Δ*ack-pta* mutant and the Δ*ack-pta*/*ack-pta*,*Δack-pta/acs+Ppase*_MTH_ complementation strains in high-salt (HS) medium supplemented with 40 mM acetate. Doubling times (T_D_) and lag times (T_Lag_) are reported in hours ± standard deviation. Significant difference between the T_D_ and T_Lag_ of various mutant strains compared to the WT was determined by calculating p-values using a two-sided t-test assuming unequal variance. If no increase in optical density at 600 nm was observed after 6 months, we assumed that there was no growth. “n.g.” = no growth.

| Substrate | Strain | $T_D$(hr)<sup>†</sup> | $T_D$ p-val | $T_{Lag}$ (hr) | $T_{Lag}$ p-val |
| --- | --- | --- | --- | --- | --- |
| 40 mM<br>Acetate | WT | 164.3 ± 5.1 | - | 498.6 ± 51.9 | - |
| | $\Delta ack-pt a$ | n.g. | - | n.g. | - |
| | $\Delta ack-pt a / ack+pt a$ | 141.2 ± 22.7 | 0.23 | 579.5 ± 80.9 | 0.2 |
| | $\Delta ack-pt a / acs+PPase_{MTH}$ | n.g. | - | n.g. | - |

<sup>†</sup> The typical doubling time for WWM60 on HS+acetate is ~80 hours. However, given that our $ack+pt a$ complementation strain demonstrates a similar doubling time, we attribute the slower growth phenotypes in this experiment to batch-specific effects in the HS media. Further, given that we do not compare or claim significance across experiments, these batch-specific effects on growth are well-controlled within this experiment.

Next, we expressed Acs homologs from *Methanothrix soehngenii*, *Methanothrix harundinacea*, and *Methanomethylovorans hollandica* in *M. acetivorans*. All the catalytically important residues and substrate-coordination motifs are conserved in the Acs sequences we selected for this study (**Fig. S3**) (21). First, to test for any toxic side-effects of Acs, we expressed these genes in WT and did not observe any atypical growth upon induction (**Table S4**). These data suggest that the expression of Acs, in and of itself, is not toxic in *M. acetivorans*. However, when we complemented the Δ*ack-pta* strain with each of these Acs homologs, no observable growth on acetate was detected, even after 6+ months of incubation (**Table S4**). Since Acs hydrolyzes ATP to AMP and pyrophosphate (PPi), we reasoned that PPi buildup might limit the growth in *M. acetivorans*. So, we generated additional constructs expressing Acs in conjunction with a pyrophosphatase (PPase) from *M. harundinacea* (**Table S4**) (22). Furthermore, we also measured acetyl-CoA production in the strain expressing *acs* and *PPase* from *M. harundinacea* (*Δack-pta/PmcrB(tetO4)-acs+PPase*_MTH_). While we could detect 10-fold higher acetyl-CoA production in the Acs-encoding cell lysates relative to the Δ*ack-pta* mutant (**Fig. 2A**), even maximum induction of Acs and PPase did not restore growth on acetate (**Fig. 2B**, **Table 1, Table S4**).

Finally, we hypothesized that the bifunctional acetyl-CoA synthase/carbon monoxide dehydrogenase enzyme (Acs/CODH), which catalyzes the dismutation of acetyl-CoA, might not be expressed as highly on TMA or in the absence of Ack+Pta on acetate. To test this hypothesis, we obtained the expression profiles of the *cdh* operons for WT on acetate using RNA-sequencing and compared it to a previously published dataset for the same strain on TMA (11). The native expression of both *cdh* operons on acetate (average log_2_(FPKM) = 9.38 for *cdh1* and average log_2_(FPKM) = 10.28 for *cdh2*) was higher than on TMA (average log_2_(FPKM) = 7.58, and 7.74 for *cdh1* and *cdh2,* respectively). The data corroborated previous reports (23), and prompted us to experimentally test if lower *cdh* expression might prevent growth on acetate in the Acs-encoding strains. Thus, we modified the plasmid encoding Acs and PPase from *M. harundinacea* to constitutively expresses the *cdh2* operon from the *Methanosarcina mazei* P*mcrB* promoter (Δ*ack-pta*/*PmcrBMM-cdh2, PmcrB(tetO1)acs-PPase_MTH_*). However, even with constitutively high expression of Acs/CODH, no growth was detected on acetate (**Table S4**).

Since nearly every Category III methanogen also encodes the Fpo complex (40/42 genomes) (**Fig. S1, Table S1**), we posited that this complex might have a role in energy conservation during acetoclastic methanogenesis. Since Rnf is likely the preferred enzyme catalyzing the re-oxidation of Fd_red_ in *M. acetivorans,* we expressed the Acs and PPase from *M. harundinacea* in the Δ*mmcA-rnf* mutant (*ΔmmcA-rnf/ PmcrB(tetO4)-acs+PPase*_MTH_). In the Δ*mmcA-rnf* mutant background, Fpo is the only known ion-translocating bioenergetic complex that can interact with Fd_red_ in the Δ*mmcA-rnf* mutant and is also upregulated by 1-to 3-fold (11). However, this mutant was also not viable on acetate (**Table S4**). Together, our results indicate that, despite being functional, Acs alone, or with a PPase, cannot replace Ack+Pta as an acetate activation module in *M. acetivorans*.

### Overexpression of Fpo’ does not restore acetoclastic growth in the Δ *mmcA-rnf* mutant

Unlike category III strains that are commonly found, a putative category IV strain, i.e., one encoding Ack + Pta and Fpo’, is yet to be discovered. This strain could, in principle, grow on acetate. *M. acetivorans* and many other strains that belong the Category II acetoclastic methanogens encode all the *fpo* genes in addition to Rnf (or Ech) (**Fig. 1**). Thus, if Fpo’ were to be made in *M. acetivorans,* it could compensate for Rnf during growth on acetate, but this likely does not happen due to regulatory constraints. Our hypothesis is based on transcriptomic analyses, which indicate that the all the *fpo* genes in *M. acetivorans* have significantly lower expression on acetate (average log_2_(FPKM) = 4.83) compared to our previously reported values on TMA (average log_2_(FPKM) = 6.45) (11). To test our hypothesis, we generated two classes of mutants that would specifically overexpress some or all of the *fpo* genes on acetate.

First, we targeted a known repressor of the *fpo* genes, *mreA* (MA3302, MA_RS17255) (24). We deleted *mreA* in the wildtype background and the Δ*mmcA-rnf* to generate Δ*mreA* and Δ*mmcA-rnf*Δ*mreA* mutants, respectively, and verified these mutants by whole genome sequencing (**Tables S2, S5**). The growth pattern of the *ΔmreA* and Δ*mmcA-rnf*Δ*mreA* mutants phenocopied their parent strains on TMA (**Fig. 3A**, **Table 2**). The Δ*mreA* mutant could still grow on acetate, albeit with a substantial fitness defect (**Fig. 3B**, **Table 2**) whereas the Δ*mmcA-rnf*Δ*mreA* strain was unable to grow even after 6+ months of incubation (**Fig. 3B**, **Table 2**). We also performed RNA-sequencing of WT and the Δ*mreA* mutant on acetate to confirm that the expression of all the *fpo* genes is, indeed, elevated when MreA is absent (**Fig. 3C**). In line with previous observations, we also observed significantly lower expression of acetate kinase (*ack*) and phosphotransacetyalase (*pta*) in the Δ*mreA* mutant (**Fig. 3D**), which could be the reason for the fitness defect observed during growth on acetate (**Fig. 3B**, **Table 2**) (24).

**Figure 3.**
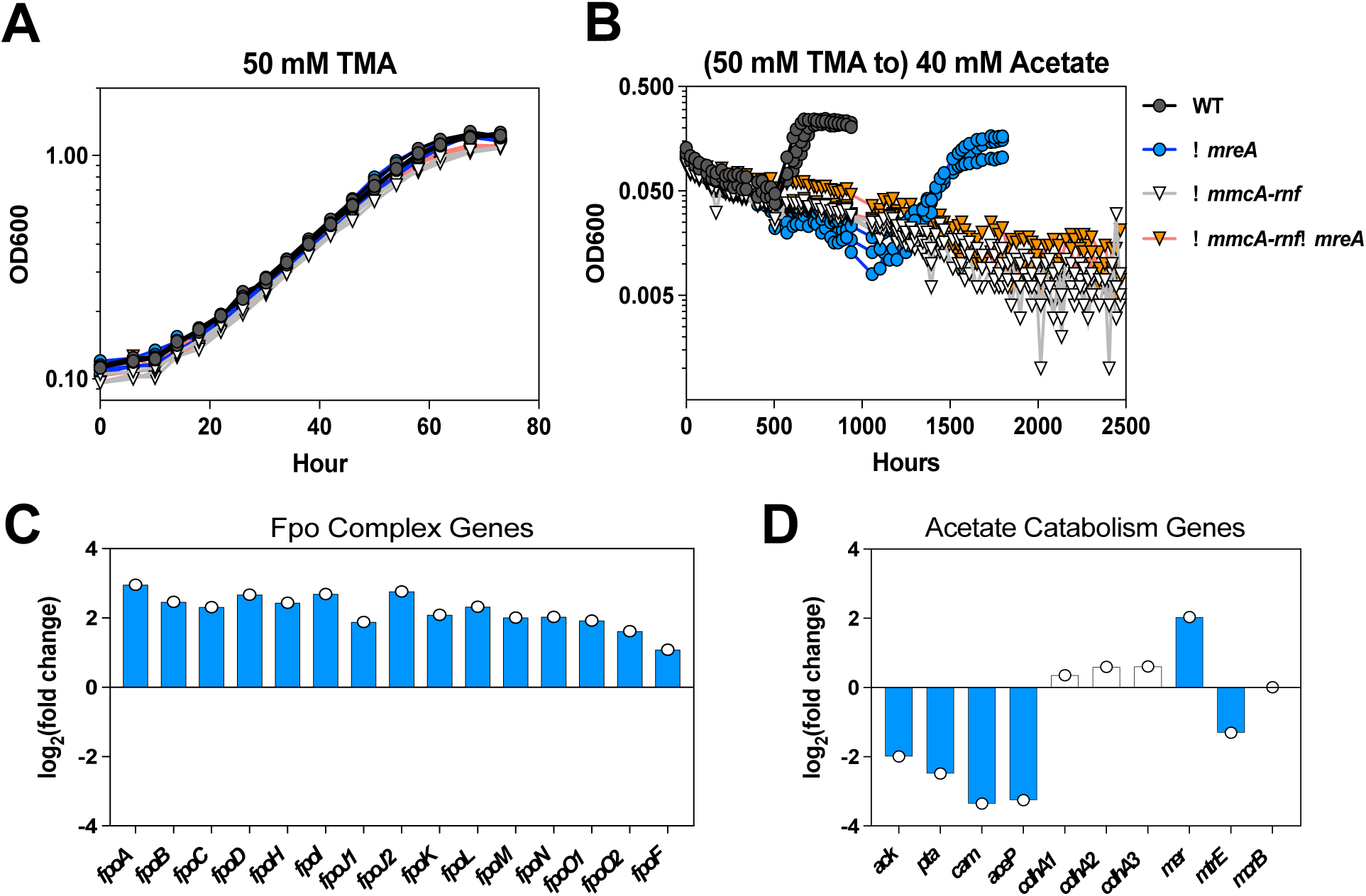
Deletion of *mreA* increases the expression of the *fpo* genes but does not permit acetate growth in the absence of the Rnf complex. Growth curve of WWM60 (wildtype or WT, black circles), Δ*mreA* (blue circles), Δ*mmcA-rnf* (white inverted triangles) and Δ*mmcA-rnf*Δ*mreA* (orange inverted triangles) strains in high-salt (HS) minimal medium with (**A**) 50 mM trimethylamine (TMA) or (**B**) 40 mM acetate after pre-culture in medium with 50 mM TMA. Four replicates were used for growth assays on TMA and three replicates were used for growth assays on acetate. (**C**) Log_2_(fold change) in expression of the *fpo* genes in the Δ*mreA* mutant compared to WT during growth on acetate. Genes with higher expression in the Δ*mreA* mutant have a positive log_2_(fold change) value. (**D**) Log_2_(fold change) in expression of metabolic genes in the Δ*mreA* mutant compared to WT during growth on acetate. For catabolic enzymes composed of more than one subunit, only expression of the first gene in the operon is shown for simplicity. Genes with higher expression in the Δ*mreA* mutant have a positive log_2_(fold change) value. Genes with lower expression in the Δ*mreA* mutant have a negative log_2_(fold change) value. Statistically significant changes in gene expression (q-value ≤ 0.01) are shown in blue bars. Statistically insignificant changes in gene expression (q-value > 0.01) are shown inwhite bars. Gene abbreviations: *fpo*, F420:phenazine oxidoreductase; *ack*, acetate kinase; *pta*, phosphotransacetylase; *cam*, carbonic anhydrase; *aceP*, acetate permease; *cdh*, carbon monoxide dehydrogenase/acetyl-CoA synthase; *mer*, methylenetetrahydromethanopterin reductase; *mtr*, H4MPT:CoM methyltransferase; *mcr*, methyl coenzyme M reductase. Growth parameters for each strain are shown in Table 2.

**Table 2.**
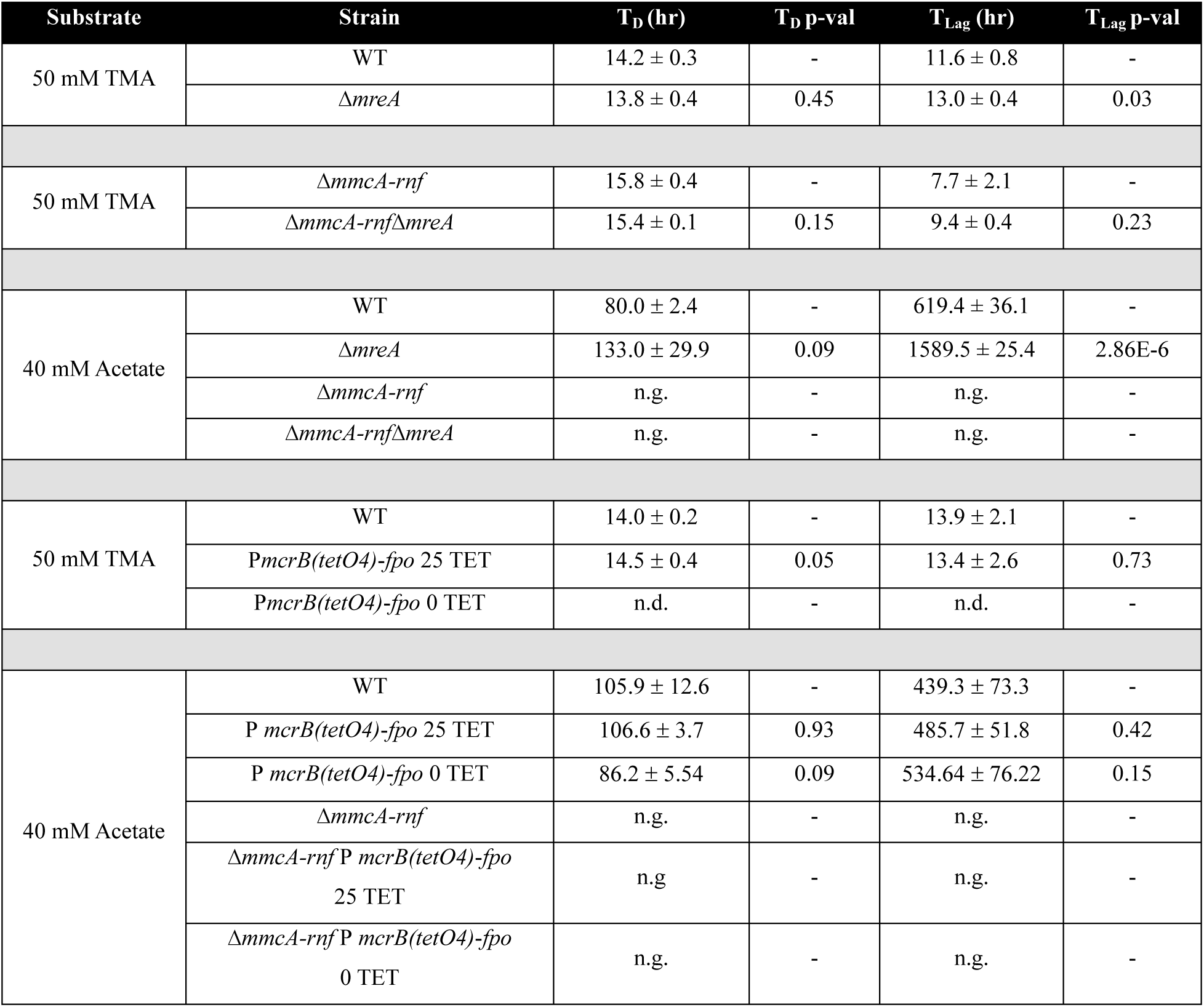
Growth characteristics of WWM60 (wildtype or WT), Δ*mreA*, Δ*mmcA-rnf*, Δ*mmcA-rnf*Δ*mreA*, P*mcrB(tetO4)-fpo*, and Δ*mmcA-rnf* P*mcrB(tetO4)-fpo* strains in high-salt (HS) minimal media with either 50 mM trimethylamine (TMA) or 40 mM acetate. . Doubling times (T_D_) and lag times (T_Lag_) are reported in hours ± standard deviation. Significant difference between the T_D_ and T_Lag_ of various mutant strains compared to the WT was determined by calculating p-values using a two-sided t-test assuming unequal variance. In each experiment, the reported p-values are determined by comparing the mutant strain against its respective parent strain and are not measured across experiments. If no increase in optical density at 600 nm was observed after 6 months, we assumed that there was no growth. “n.g.” = no growth. T_D_ and T_Lag_ were not determined for strains/conditions that did not grow exponentially. “n.d.” = not determined.

Since the deletion of MreA downregulates acetate activation genes and leads to a global transcriptional response (**Fig. S4, Table S6**) (24), we designed an alternate strategy for targeted overexpression of Fpo’ in *M. acetivorans*. The Fpo complex is expected to contain 13 subunits, 12 of which are encoded as a single ∼10.5 kb operon, *fpoABCDHJ_1_J_2_KLMNO_1_* (MA1495-1507, MA_RS07760-07820) (18). An additional FpoO subunit, *fpoO2* (MA1509, MA_RS25205), is encoded nearby but its role is undefined (18). One additional subunit, FpoF (MA3732, MA_RS19445), the F_420_ interacting “head” of the complex, is found elsewhere in the genome (18). In obligately acetoclastic methanogens the *fpoF* locus is absent, which likely enables the rest of the Fpo complex to operate as a ferredoxin: methanophenazine oxidoreductase (14). Accordingly, we hypothesized that increasing expression of the *fpoA-O_1_*operon, without changing the expression of the *fpoF* gene, might generate more of the Fpo’ complex that could interface with Fd_red_ produced during acetoclastic growth. To this end, we replaced the native promoter of the *fpo* operon with a tetracycline-inducible promoter [P*mcrB(tetO4*)] in both WT [P*mcrB(tetO4*)-*fpo*] and the Δ*mmcA-rnf* mutant [Δ*mmcA-rnf* P*mcrB(tetO4*)-*fpo*] (**Fig. S5A, S5B**, **Tables S2, S5**). To validate tetracycline-inducible control of the *fpoA-O*_1_ operon, we grew the P*mcrB(tetO4*)-*fpo* strain in varying concentrations of tetracycline and observed linear growth at 0 µg/mL tetracycline indicating that the mutant becomes limited for Fpo (25, 26), which is known to be essential for growth on TMA (**Fig. 4A**, **Table 2**) (8, 12). In the Δ*mmcA-rnf* P*mcrB(tetO4*)-*fpo* mutant, which lacks the TetR repressor, the promoter swap results in constitutive expression of the *fpoA-O_1_*operon and therefore does not limit growth on TMA (**Fig. S6, Table S7**). When we transferred these mutants to acetate, the P*mcrB(tetO4*)-*fpo* strain could still grow but the Δ*mmcA-rnf* P*mcrB(tetO4*)-*fpo* mutant was incapable of growth after 6+ months of incubation (**Fig. 4B**, **Table 2**). Overall, our results indicate that increasing the expression of the Fpo’ complex in *M. acetivorans* cannot compensate for the Rnf complex during growth on acetate.

**Figure 4.**
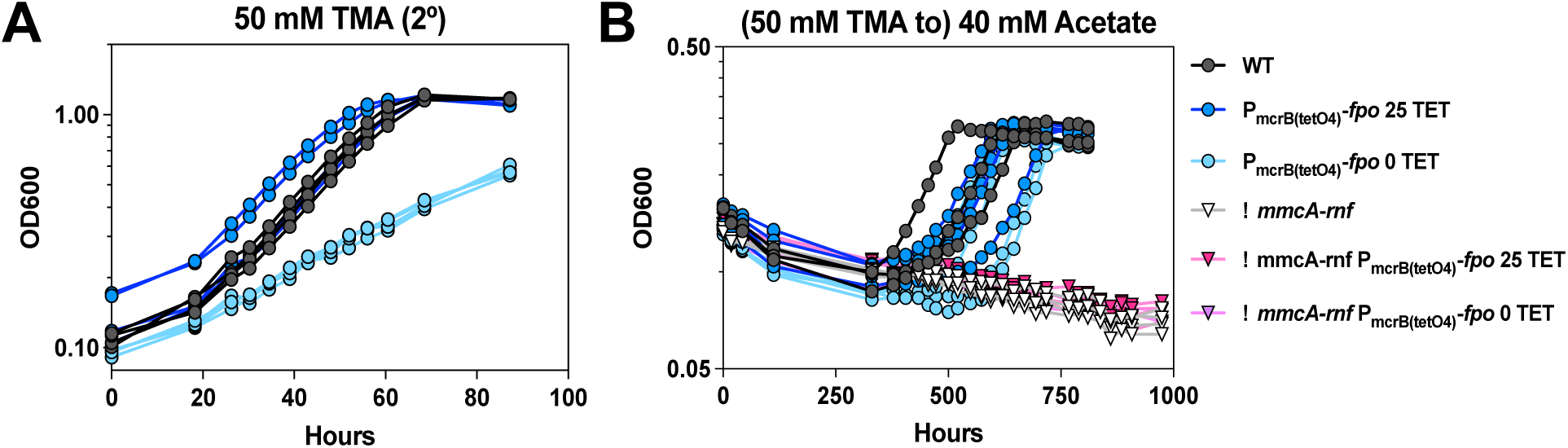
Overexpression of the *fpo’* genes does not rescue acetoclastic growth of the Δ*mmcA-rnf* mutant. (**A**) Growth curves of WWM60 (wildtype or WT, black circles) and the P*mcrB(tetO4)-fpo* mutant (blue circles) in high-salt (HS) minimal medium with 50 mM trimethylamine (TMA). The P*mcrB(tetO4)-fpo* mutant is supplemented with 25 µg/mL tetracycline (dark blue circles) or 0 µg/mL tetracycline (light blue circles). Growth assays were conducted with cultures pre-grown in HS with 50 mM TMA with the aforementioned concentrations of tetracycline (**B**) Growth of WT (black circles), the P*mcrB(tetO4)-fpo* mutant strain supplemented with 25 µg/mL tetracycline (dark blue circles) or 0 µg/mL tetracycline (light blue circles), Δ*mmcA-rnf* (white triangles), and Δ*mmcA-rnf* P*mcrB(tetO4)-fpo* mutant strains provided with 25 µg/mL tetracycline (pink triangles) or 0 µg/mL tetracycline (purple triangles) in HS minimal medium with 40 mM acetate after pre-culture in HS medium with 50 mM TMA. Four replicate tubes for each strain were used for growth assays. Note: One replicate for WT broke during the experiment and all data for that replicate has been omitted. Growth parameters for each strain are shown in Table 2.

## DISCUSSION

Our bioinformatic screen of methanogens within the Class *Methanosarcinia* suggested that the genomic potential for acetoclastic methanogenesis is broader in scope than the two Genera where this metabolism has been previously demonstrated (**Fig. 1D**). Although no methanogens that encode Acs and Rnf, i.e., the Category III strains, have been shown to grow on acetate as the sole carbon and energy substrate, we decided to pursue a more formal test of this hypothesis using *M. acetivorans*. We reasoned that this direct genetic approach was more appropriate than re-attempting cultivation experiments with Category III strains, as there is a wealth of experimental data describing how *M. acetivorans* grows on acetate, which allowed us to test the role of individual modules in the acetoclastic pathway.

To explore acetate metabolism via Acs and Rnf, we generated a Δ*ack-pta* strain of *M. acetivorans*, which cannot grow on acetate due to a disruption in the substrate activation pathway (**Fig. 2**). Our data validates previous reports that transposon insertions into these genes abrogates growth on acetate (27). However, when we attempted to complement our Δ*ack-pta* mutant with *acs* sequences from across the *Methanosarcinia*, these genes failed to rescue acetoclastic growth despite measurable acetyl-CoA production in cell lysates. We note reduced acetyl-CoA production in the *Δack-pta/acs* mutants compared to the *Δack-pta/ack-pta* complementation strain but interpret this outcome to be reflective of the lower V_max_ of Acs vs Ack, as has been previously reported (5, 19). Further, reduced acetyl-CoA production would only slow down growth of *Δack-pta/acs* mutants compared to WT, which would be more in line with previous observations of slower growth rates among the *Methanothrix* spp. compared to the *Methanosarcina* spp. (5). The complete lack of growth suggests that the Acs and Rnf modules are functionally incompatible for acetoclastic growth in methanogens. Based on this outcome, we hypothesize that Category III methanogens might retain Acs for the purpose of biomass production from acetate, which could be confirmed with future work in these strains.

In parallel, we also addressed the likelihood of acetate growth in a putative Class IV strain, i.e., one that would encode Ack+Pta and Fpo’. Although this combination of genes was not detected in any genome in our bioinformatic screen, Fpo’ is hypothesized to be the energy conservation module required for acetoclastic growth in *Methanothrix*, but this has not been directly tested due to the genetic intractability of *Methanothrix* spp. (14, 15). Additionally, mutants of *Methanosarcina mazei* lacking Ech, can still generate a chemiosmotic gradient with Fd_red_, which is hypothesized to be mediated by the Fpo’ complex (28). Therefore, we used mutants of *M. acetivorans* that express the Fpo’ complex instead. Our results suggest that there is a fundamental incompatibility between the Ack+Pta and Fpo’ as modules for mediating acetoclastic growth. One plausible reason is that the presence of FpoF in *M. acetivorans* precludes the rest of the Fpo complex, i.e., Fpo’, from interacting with Fd. Alternately, there might also be incompatibility between the Fd_red_ produced during acetate metabolism and the active site of the potential Fpo’ complex generated in our mutants, as ferredoxin specificity has been noted for other metabolic processes in bacteria and archaea (29–33). It should also be noted that despite encoding Acs and Fpo’, *Methermicoccus shengliensis* has not been reported to grow with acetate as the sole substrate (34, 35). Thus, the lack of acetoclastic growth in our Δ*mmcA-rnf* P*mcrB*(*tetO4*)-*fpo’* strain might be due to additional proteins in *Methanothrix* spp. strains that modify the Fpo’ to interact with Fd, as has been observed for Fd-interacting Complex I homologs in chloroplasts (36, 37). Future work on the *Methanothrix* Fpo’ complex will help to answer refine how this complex interacts with Fd.

Our results point to incompatibility between modules of acetate activation and energy conservation that have not already been described in isolated acetoclastic methanogens (**Fig. 5A**) and therefore highlight an important nuance in the evolutionary history of acetoclastic methanogenesis within the *Methanosarcinia*. Previous hypotheses have implicated horizontal gene transfer as being the major driver of acetoclastic metabolism among Ack+Pta containing methanogens (6, 38), but our results emphasize that the horizontal acquisition of catabolic genes alone is not sufficient, and that integration with energy-conserving modules of the ETC would also be required. Thus, the transitions to acetoclastic metabolism that have occurred among the common ancestors of the extant *Methanosarcinia* have been shaped by coordinated evolutionary trajectories of the ETC among the *Methanothrix*, and the substrate activation genes among the *Methanosarcina* (**Fig. 5B**). Building on this study, future work is necessary for detailing the physiological and ecological adaptations that facilitate acetoclastic methanogenesis in *Methanosarcina* spp. and *Methanothrix* spp.

**Figure 5.**
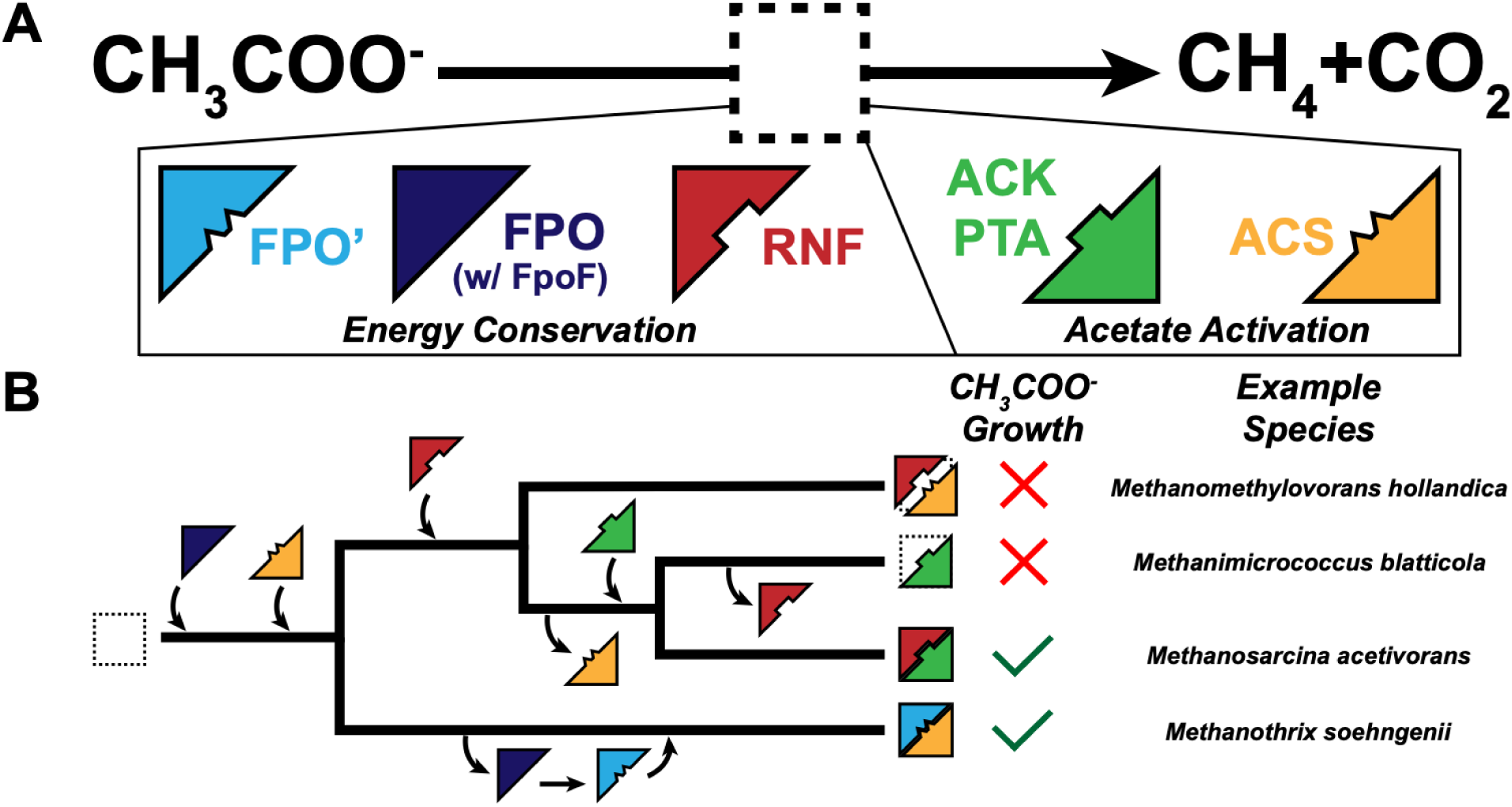
Co-evolution of energy conservation and acetate activation modules is necessary to enable acetoclastic methanogenesis. (**A**) Model depicting energy conservation and acetate activation modules as colored blocks that must interact precisely to fill in the empty square representing acetoclastic methanogenesis. (**B**) The provenance of enzyme modules from Panel A among select methanogens within the Class *Methanosarcinia*. An ancestral strain might have encoded Fpo (dark blue) along with Acs (yellow). However, to enable acetoclastic growth in the *Methanothrix*, Fpo would have had to undergone loss of FpoF and potentially further modification to generate Fpo’ (light blue) compatible with Acs for acetoclastic methanogenesis. In the other methanogens, Rnf (red) was likely acquired after the most recent common ancestor of Category I diverged from the *Methanotrichales*. Methanogenic ancestors with Rnf were likely precluded from using acetate as a sole substrate until Acs was replaced with Ack+Pta (green) in the ancestor of *Methanosarcina* and *Methanomicrococcus* facilitated a new form of acetoclastic methanogenesis (Category II). Loss of Rnf in *Methanomicrococcus* due to their adaptation to host-associated niches led to a loss of acetoclastic methanogenesis in this group of methanogens.

## ACKNOWLEDGEMENTS

We thank all members of the Nayak lab for their valuable feedback and support. We would like to especially thank Annelise Goldman for technical assistance with growth curves. DDN acknowledges funding from the Searle Scholars Program sponsored by the Kinship Foundation, the Rose Hills Innovator Grant, the Beckman Young Investigator Award sponsored by the Arnold and Mabel Beckman Foundation, the Alfred P. Sloan Research Fellowship sponsored by the Sloan Foundation, the Simons Foundation Early Career Investigator in Marine Microbial Ecology and Evolution Award, and the Packard Fellowship in Science and Engineering sponsored by the David and Lucille Packard Foundation. DDN is a Chan-Zuckerberg Biohub – San Francisco Investigator. KES was supported in part by the NSF Graduate Research Fellowship Program (Fellow ID: 202299857). DG acknowledges support from the National Institute of General Medical Sciences of the National Institutes of Health for a Kirschstein-NRSA postdoctoral fellowship (F32GM150233). DDN, BED acknowledge funding from the Department of Energy through project number S589706. The funders had no role in the conceptualization and writing of this manuscript or the decision to submit the work for publication.

## DATA AVAILABILITY STATEMENT

Sequencing data have been deposited in the Sequencing Reads Archive and the BioProject number will be made available upon publication. All other data generated in this study are provided in the manuscript.

## DECLARATION OF INTERESTS

The authors declare no competing interests.

## MATERIALS AND METHODS

### Survey of Genome Taxonomy Database (GTDB)

Genomes from GTDB r214.0 (https://data.gtdb.ecogenomic.org/releases/release214/214.0/) were downloaded and annotated with Prokka v.1.14.6 using the prokaryotic genetic code 11 (39). Genome accessions belonging to the Class *Methanosarcinia* were determined using GTDB taxonomy metadata and the associated annotated genomes were selected for downstream genomic surveys. Note that the species we referred to as “*Methanothrix harundinacea*” in our analyses are currently being scrutinized for reclassification as “*Methanocrinis harundinaceus*” as proposed by Khomyakova, et al. (40). However, since the reclassification has not been formalized, we chose to refer to these organisms using their previous designation within the “*Methanothrix*” Genus. CheckM completeness scores were calculated using a concatenated biomarker set described in (https://data.gtdb.ecogenomic.org/releases/release214/214.0/METHODS.txt) (41). We obtained Profile HMMs for acetate catabolism: acetate kinase (Ack), phosphotransacetylase (Pta), and AMP-forming acetyl-CoA synthetase (Acs), and respiratory complexes: the *<u>R</u>hodobacter* <u>n</u>itrogen fixation complex (in methanogens MmcA-RnfCDGEAB), the <u>E</u>nergy <u>c</u>onverting <u>h</u>ydrogenase complex (EchABCDEF), and the <u>F</u>_420_:<u>p</u>henazine <u>o</u>xidoreductase (FpoABCDHIJKLMNOF) from the NCBI or KoalaFAM (42). A list of all accessions used is available in **Table S8**. We used a custom command line tool (https://github.com/kshalv/hmm_tools/tree/main, (43) to generate distributions for proteins involved in the activation of acetate to acetyl-CoA and energy conservation . Briefly, we used the ‘hmm’ option of the command line tool to iterate through the given profile HMM’s with HMMER 3.4 (hmmer.org, (44)), and recorded hits with an e-value threshold lower than a 1E-03 threshold. In our initial analyses, we used the trusted cutoff (TC) score threshold for all queries. We noticed that genomes belonging to the family *Methanotrichaceae* were lacking hits for several of the subunits of the Fpo. We inspected complete genomes of *Methanothrix soehngenii* (NCBI Accession: NC_015416.1) and *Methanothrix harundinacea* (NCBI Accession: NC_017527.1) and were able to identify missing subunits manually. Thus, to capture the potential diversity of Fpo complexes across all *Methanosarcinia*, we ran our HMM analysis again using the noise cutoff (NC) score threshold for all Profile HMMs. We found that the NC threshold recapitulated the TC threshold results for all proteins except Fpo, in which more subunits were identified among the *Methanotrichaceae*. Thus, we reasoned that the NC hit score prediction was accurate to cellular physiology. All protein hits which exceeded the noise cutoff (NC) threshold were counted and recorded for each genome to generate a presence/absence distribution table. The presence/absence hit distribution was overlaid on a tree of *Methanosarcinia* that was generated from the ar53_r214.tree available through GTDB. For energy conservation complexes composed of multiple subunits, a threshold number of hits had to be successfully detected for “presence” to be counted: For RNF, ≥ 4 of 6 subunits; for FPO’, ≥ 5 of 7 subunits; for ECH, ≥ 4 of 6 subunits.

### Media and Culture Conditions

*M. acetivorans* strains were grown in bicarbonate-buffered high-salt (HS) liquid medium containing either 50 mM TMA, 40 mM sodium acetate, or a combination of 50 mM TMA and 20 mM acetate as the growth substrate and an 80:20% mix of N_2_:CO_2_ gas in the headspace at approximately 55-69 kPa (45). All substrates were added prior to autoclaving. For all liquid culturing, *M. acetivorans* strains were incubated without shaking at 37 °C. To generate mutants, cells were plated on 50 mM TMA with 1.5% w/v agar (Sigma-Aldrich, St. Louis, MO, USA) also containing 2 µg/mL Puromycin as a selective agent. Puromycin was added to agar-solidified HS-TMA after autoclaving from a 1000X sterile, anaerobic stock solution with N_2_ gas in the headspace at approximately 55-69 kPa. Agar-solidified HS-TMA plates were incubated at 37 °C in a custom-built inter-chamber anaerobic incubator with a headspace of 79.9:20:0.1% N_2_:CO_2_:H_2_S, as described previously (46). Where indicated, mutant strains were cultured with 2 µg/mL puromycin to maintain plasmids or various amounts of tetracycline hydrochloride to induce expression of genes from tetracycline-inducible promoters as described (47). Puromycin and tetracycline were added to culture tubes after autoclaving but before inoculation from 10X or 100X filter-sterilized, anaerobic stock solutions with N_2_ gas in the headspace at 55-69 kPa. *Escherichia coli* strains were routinely grown using lysogeny broth (LB). For liquid culturing, strains were incubated at 37 °C shaking at 250 rpm in a shaking incubator (Thermo Fisher Scientific, Waltham, MA, USA). For mutant generation, strains were plated on LB with 1.5% w/v agar and antibiotics as a selective agent: 25 µg/mL kanamycin and/or 10 µg/mL chloramphenicol. For plasmid extraction, strains were grown in liquid LB with equivalent concentrations of antibiotics as agar plates and 10 mM rhamnose.

### Plasmid Construction

For CRISPR-Cas9 mediated gene editing, plasmids were constructed as described previously (48) using Gibson assembly (49). Briefly, 20 bp guide sequences were designed against the *M. acetivorans* C2A genome using the Find CRISPR Sites tool with an NGG 3’ PAM site using the Geneious platform (v.11.0). The 20-bp guides were synthesized as overhangs on primers (Integrated DNA Technologies, Coralville, IA, USA) used to introduce the sgRNA cassette into *AscI*-digested pDN201 (48). A list of sgRNA targeting sequences can be found in **Table S9**. In the same plasmid backbone at the *PmeI* site, a *ca.* 2kbp homology directed repair template was cloned in to generate the edits of interest at the sites cut by the sgRNA cassette. For in-frame gene deletions, *ca.* 1kb upstream and *ca.* 1kb downstream of the gene(s) of interest were amplified from the chromosome, leaving only 30bp of the 5’ and 3’ ends of the gene(s) in the repair template. For the promoter swap mutations, a commercial DNA fragment (Integrated DNA Technologies, Coralville, IA, USA) encoding the desired terminator and tetracycline-inducible P*mcrB(tetO4)* promoter (∼400bp) was fused with *ca.* 850 bp upstream and *ca.* 730 bp downstream homology arms amplified via PCR from the chromosome. Complementation plasmids were also generated via Gibson assembly (49) by cloning in the gene(s) of interest either from commercial gene fragments (Twist Bioscience, South San Francisco, CA, USA) or from genes amplified via PCR from the chromosome of *M. acetivorans* or *M. hollandica* under a tetracycline-inducible P*mcrB(tetO1)* promoter in pJK027A (47). The pDN201- and pJK027A-derived plasmids were retrofitted with pAMG40 for autonomous replication in M. acetivorans using Invitrogen Gateway BP Clonase II Enzyme mix (Thermo Fisher Scientific, Waltham, MA, USA) as previously described (47). Plasmids were transformed into *E. coli* strain WM4489 by electroporation (MicroPulser Electroporator, Bio-Rad, Hercules, CA, USA). Plasmids were extracted from host *E. coli* strains using the Zymo Zyppy Plasmid Miniprep Kid (Zymo Research, Irvine, CA, USA). Plasmids were confirmed using PCR and Sanger sequencing (Barker Sequencing Facility, UC Berkeley, Berkeley, CA, USA) or restriction endonuclease digestion. Plasmids used for mutant generation are listed in **Table S10**. A complete list of primers used in this study is available in **Table S11**.

### *M. acetivorans* Mutant Generation

Mutants of *M. acetivorans* were generated using a liposome-mediated transformation protocol inside an anaerobic chamber with a gas atmosphere of 78:18:4% N_2_:CO_2_:H_2_ as previously described (50). In brief, 20 mL of late exponential phase cultures (OD600 ∼ 0.8-1.0) were harvested by centrifugation in the anaerobic chamber. The cell pellet was resuspended in 750 µL of anaerobic, isotonic, bicarbonate-buffered sucrose (pH = 7.4) containing 100 µM cysteine. Next, 25 μL of N-[1-(2,3-Dioleoyloxy)propyl]-N,N,N-trimethylammonium methylsulfate (DOTAP, Roche Diagnostics Deutschland GmbH, Mannheim, Germany) or 3 µL of DOTAP Chloride (MedChemExpress, Monmouth Junction, NJ, USA) and 2 μg of plasmid DNA resuspended in 50 µL of sucrose buffer were pre-incubated with 75 µL (or 97 µL) of anaerobic buffered sucrose for 30 minutes before being added to the cell suspension mixture. Transfections with the DOTAP+DNA liposomes and cells were incubated at room temperature for 4 hours in the anaerobic chamber before inoculation into 10 mL of 50 mM HS-TMA. Outgrowths of transfected cells were incubated at 37 °C for 12-16 hours before plating on 50 mM HS-TMA agar solidified medium with 2 µg/mL Puromycin as a selective agent. For clearing the CRISPR-Cas9-containing plasmids from deletion mutants, strains were plated on 50 mM HS-TMA agar with 20-160 µg/mL 8-aza-diaminopurine (8ADP). Strains were genotyped for clearance of the mutagenic plasmid by diagnostic PCR for *pac*, the puromycin resistance gene, and/or whole genome sequencing. A full list of strains used in this study is available in **Table 3**.

**Table 3.** List of *M. acetivorans* strains used in this study.

| Strain | Genotype | Construction Details | Source |
| --- | --- | --- | --- |
| WWM60 | $\Delta hpt::P_{mcrB}\text{-}tetR$ | - | Guss, et al. (2008) |
| WWM1015 | $\Delta hpt::P_{mcrB}\text{-}tetR \Delta mmcA\text{-}rnf$ | - | Mand, (2018) |
| DDN227 | $\Delta hpt::P_{mcrB}\text{-}tetR \Delta mreA$ | WWM60 was transformed to Pur<sup>R</sup> with pBD014; plasmid-cured strain was isolated by plating on medium with 8ADP | This study |
| DDN230 | $\Delta hpt::P_{mcrB}\text{-}tetR \Delta mmcA\text{-}rnf \Delta mreA$ | WWM1015 was transformed to Pur<sup>R</sup> with pBD014; plasmid-cured strain was isolated by plating on medium with 8ADP | This study |
| DDN235 | $\Delta hpt::P_{mcrB}\text{-}tetR \Delta ack\text{-}pta$ | WWM60 was transformed to Pur<sup>R</sup> with pBD024; plasmid-cured strain was isolated by plating on medium with 8ADP | This study |
| DDN264 | $\Delta hpt::P_{mcrB}\text{-}tetR T_{mcr} P_{mcrB}(tetO4)\text{-}fpo$ | WWM60 was transformed to Pur<sup>R</sup> with pBD030; plasmid-cured strain was isolated by plating on medium with 8ADP | This study |
| DDN306 | $\Delta hpt::P_{mcrB}\text{-}tetR \Delta mmcA\text{-}rnf T_{mcr} P_{mcrB}(tetO4)\text{-}fpo$ | WWM60 was transformed to Pur<sup>R</sup> with pBD030; plasmid-cured strain was isolated by plating on medium with 8ADP | This study |
| DDN345 | WWM60/pBD035<br>[$P_{mcrB}(tetO1)\text{-}acs3_{MTS}$] | WWM60 was transformed to Pur<sup>R</sup> with pBD035; isolates were verified by Sanger sequencing and grown in HS medium supplemented with 2 µg/mL Puromycin | This study |
| DDN347 | WWM60/pBD044<br>[$P_{mcrB}(tetO1)\text{-}acs3_{MMH}$] | WWM60 was transformed to Pur<sup>R</sup> with pBD044; isolates were verified by Sanger sequencing and grown in HS medium supplemented with 2 µg/mL Puromycin | This study |
| DDN348 | DDN235/pBD043<br>[$P_{mcrB}(tetO1)\text{-}ack\text{+}pta$] | DDN235 was transformed to Pur<sup>R</sup> with pBD043; isolates were verified by Sanger sequencing and grown in HS medium supplemented with 2 µg/mL Puromycin | This study |
| DDN349 | DDN235/pBD044<br>[$P_{mcrB}(tetO1)\text{-}acs3_{MMH}$] | DDN235 was transformed to Pur<sup>R</sup> with pBD044; isolates were verified by Sanger sequencing and grown in HS medium supplemented with 2 µg/mL Puromycin | This study |
|  |  | Puromycin |  |
| DDN350 | WWM60/pBD037<br>[*P<sub>mcrB</sub>(tetO1)-acs3<sub>MTH</sub>*] | WWM60 was transformed to Pur<sup>R</sup> with pBD037; isolates were verified by Sanger sequencing and grown in HS medium supplemented with 2 µg/mL Puromycin | This study |
| DDN352 | DDN235/pBD035<br>[*P<sub>mcrB</sub>(tetO1)-acs3<sub>MTS</sub>*] | DDN235 was transformed to Pur<sup>R</sup> with pBD035; isolates were verified by Sanger sequencing and grown in HS medium supplemented with 2 µg/mL Puromycin | This study |
| DDN353 | DDN235/pBD037<br>[*P<sub>mcrB</sub>(tetO1)-acs3<sub>MTH</sub>*] | DDN235 was transformed to Pur<sup>R</sup> with pBD037; isolates were verified by Sanger sequencing and grown in HS medium supplemented with 2 µg/mL Puromycin | This study |
| DDN354 | DDN235/pBD038<br>[*P<sub>mcrB</sub>(tetO1)-acs3<sub>MTH</sub>+PPase<sub>MTH</sub>*] | DDN235 was transformed to Pur<sup>R</sup> with pBD038; isolates were verified by Sanger sequencing and grown in HS medium supplemented with 2 µg/mL Puromycin | This study |
| DDN360 | WWM60/pBD036<br>[*P<sub>mcrB</sub>(tetO1)-acs3<sub>MTS</sub>+PPase<sub>MTS</sub>*] | WWM60 was transformed to Pur<sup>R</sup> with pBD036; isolates were verified by Sanger sequencing and grown in HS medium supplemented with 2 µg/mL Puromycin | This study |
| DDN361 | WWM60/pBD038<br>[*P<sub>mcrB</sub>(tetO1)-acs3<sub>MTH</sub>+PPase<sub>MTH</sub>*] | WWM60 was transformed to Pur<sup>R</sup> with pBD038; isolates were verified by Sanger sequencing and grown in HS medium supplemented with 2 µg/mL Puromycin | This study |
| DDN362 | DDN235/pBD036<br>[*P<sub>mcrB</sub>(tetO1)-acs3<sub>MTS</sub>+PPase<sub>MTS</sub>*] | DDN235 was transformed to Pur<sup>R</sup> with pBD036; isolates were verified by Sanger sequencing and grown in HS medium supplemented with 2 µg/mL Puromycin | This study |
| DDN448 | DDN235/pBD047 [*P<sub>mcrBMM</sub><sup>-</sup>cdh2, P<sub>mcrB</sub>(tetO1)-acs/PPase<sub>MTH</sub>*] | DDN235 transformed to Pur<sup>R</sup> with pBD047; isolates were verified by Sanger sequencing and grown in HS medium supplemented with 2 µg/mL Puromycin | This study |

### Genomic DNA Extraction, Sequencing, and Analysis

Approximately 2 mL of cells were harvested from a saturated culture of an *M. acetivorans* strain and genomic DNA was extracted using the Qiagen DNeasy Blood & Tissue kit (Qiagen, Hilden, Germany) according to the manufacturer’s instructions. Samples were sent for library preparation and whole genome sequencing at SeqCenter (Pittsburgh, PA, USA). Analysis of the sequencing results was conducted using *breseq* v0.35.5 (51). Sequencing reads have been uploaded to the Sequencing Reads Archive (SRA) and will be made available upon publication.

### Acetylhydroxamate Assay for Measuring Acetyl-CoA Production

A colorimetric assay to detect the production of acetyl groups from acetate in cleared cell lysates was developed based on previous methods (19, 20). Briefly, ∼30 mL of TMA-grown or ∼250-500 mL of acetate-grown *M. acetivorans* strains were harvested by centrifugation at 5000 RPM at 4°C for 15 minutes (Sorvall Legend XTR,472 Thermo Fisher Scientific, Waltham, MA, USA). Cell pellets were lysed using 600 µL of 50 mM sodium phosphate buffer (pH = 8.0) on ice. 1-2 µL of DNase was added and the lysate was incubated at room temperature for 10 mins to digest DNA. After incubation, 36 µL of 5 M sodium chloride was added to lysates and incubated for 2 minutes at room temperature. The cell lysate mixture was clarified by centrifugation at 14,000 RPM at 4 °C for 10 minutes (Sorvall Legend XTR,472 Thermo Fisher Scientific, Waltham, MA, USA). This cleared lysate was used for assays. Total protein concentration was measured in a microplate reader (BioTek Epoch 2, Winooski, VT, USA) by adding 5 µL of cleared lysate to 250 µL of Pierce Bradford Plus Protein Assay Reagent (Thermo Fisher Scientific, Waltham, MA, USA) and calibrating against a standard curve of BSA (52). The assay buffer contained 100 mM Tris HCl (pH = 8.5), 10 mM sodium acetate, 0.2 mM coenzyme A, 4 mM MgCl_2_, 2 mM adenosine 5’triphosphate disodium salt (ATP), 2 mM dithiothreitol (DTT), and 600 mM dilute, neutralized hydroxylamine. Dilute, neutralized hydroxylamine was prepared by combining equal parts of 4 M hydroxylamine HCl and 14.7% w/v potassium hydroxide which was then diluted 1:10 with MilliQ water. Reactions were started by adding 1 mg of cleared lysate to 333 µL of reaction buffer and samples were incubated at 37 °C for 30 minutes in technical duplicate or triplicate. Reactions were quenched by bringing the total reaction volume to 666 µL using 10% trichloroacetic acid to precipitate protein. Finally, 333 µL of 2.5% w/v Fe(III)Cl_3_ in 2 M HCl was added to each reaction to develop the color. Samples were measured for absorbance at 540 nm in 10x4x45 mm plastic cuvettes using a UV-Vis spectrophotometer (Genesys 50, Thermo Fisher Scientific, Waltham, MA, USA) with cuvette attachment. Absorbance of experimental samples was blanked against a no cell control reaction containing 333 µL of reaction buffer, 333 µL of 10% trichloroacetic acid, and 333 µL of Fe(III)Cl_3_. An acetyl-CoA standard curve was generated by measuring absorbance at 540 nm of cell-free reactions that contained a final concentration of 0.25, 0.5, 1, 2.5, 5, 7.5, 10, and 20 mM of acetyl-CoA (Cayman Chemical, Ann Arbor, MI, USA).

### Growth Assays

For growth analysis, cultures of *M. acetivorans* strains were inoculated into 26 mL Balch tubes containing 10 mL of fresh media, and growth was monitored as the increase in absorbance at 600 nm over time (optical density, OD600) using a UV-Vis spectrophotometer (Genesys 50, Thermo Fisher Scientific, Waltham, MA, USA). Triplicate or quadruplicate replicate tubes were used for experiments as indicated. All strains were pre-cultured in 50 mM TMA before transfer for growth analysis. For 50 mM TMA growth curves, a ∼1:20 dilution of inoculum from an early stationary phase culture was used (∼0.5 mL into 10 mL fresh medium). For 40 mM acetate growth curves, an ∼1:11 dilution of inoculum from an early stationary phase culture was used (∼1 mL into 10 mL fresh medium). Growth rates and doubling times (T_D_) were calculated from the slopes of log10-transformed OD600 values from the exponential phase by linear regression analysis with R^2^ values ≥ 0.95. Lag/acclimation times (T_Lag_) were calculated as the time in hours where the slope of the linear regression intersected with the log10-transformed OD600 value from t=0 hours (y-intercept). Analysis of growth data and statistical tests were performed in Microsoft Excel (v16.96.1). Growth curves were plotted using GraphPad Prism (v10.4.2).

### RNA Extraction, Sequencing, and Analysis

Three replicate 11 mL cultures of *M. acetivorans* strains growing on 40 mM acetate were harvested and lysed at mid-exponential phase (OD600 ∼0.100) by 1:1 addition (11 mL) of Trizol reagent (Life Technologies, Carlsbad, CA, USA) and incubated at room temperature for 5 minutes. To this mixture, 22 mL of 100% ethanol was added. RNA extraction was performed using the Qiagen RNEasy Mini Kit (Qiagen, Hilden, Germany) according to the manufacturer’s instructions. Quantification of the RNA yield was performed using a Nanodrop One/OneC UV Spectrophotometer (Thermo Fisher Scientific, Waltham, MA, USA) before storage at -80°C. Samples were shipped on dry ice to SeqCenter (Pittsburgh, PA, USA) for rRNA depletion, library preparation and Illumina sequencing. Non-interleaved, paired-end reads were uploaded to KBase, as previously described (11, 53). Reads were aligned to the *M. acetivorans* C2A genome using Bowtie (v2.3.2), assembled using Cufflinks (v.2.2.1), and differential expression analysis was performed using DESeq2 (v1.20.0). Changes in transcript abundance were determined to be “significant” if the q-value ≤ 0.01. Volcano plot for global transcriptome response and bar graphs for individual gene expression changes was generated using GraphPad Prism (v10.4.2). Sequencing reads have been uploaded to the SRA and will be made available upon publication.

